# Human-specific *ARHGAP11B* is necessary and sufficient for human-type basal progenitor levels in primate brain organoids

**DOI:** 10.1101/2020.10.01.322792

**Authors:** Jan Fischer, Jula Peters, Takashi Namba, Wieland B. Huttner, Michael Heide

## Abstract

Based on studies in various animal models, including developing ferret neocortex (Kalebic et al., 2018), the human-specific gene *ARHGAP11B* has been implicated in human neocortex expansion. However, the extent of its contribution to this expansion during primate evolution is unknown. Here we addressed this issue by genetic manipulation of ARHGAP11B levels and function in chimpanzee and human cerebral organoids. Interference with ARHGAP11B’s function in human cerebral organoids caused a massive decrease, down to a chimpanzee level, in the proliferation and abundance of basal progenitors, the progenitors thought to have a key role in neocortex expansion. Conversely, *ARHGAP11B* expression in chimpanzee cerebral organoids resulted in a doubling of cycling basal progenitors. Taken together, our findings demonstrate that *ARHGAP11B* is necessary and sufficient to maintain the elevated basal progenitor levels that characterize the fetal human neocortex, suggesting that this human-specific gene was a major contributor to neocortex expansion during human evolution.

## Introduction

The neocortex, the evolutionarily youngest part of the brain, is the seat of our higher cognitive abilities. One approach to elucidate neocortical performance has been to investigate the development of the neocortex, which has provided pivotal insight (Debra L. Silver, 2019; Dehay et al., 2015; Florio & Huttner, 2014; Lui et al., 2011; Molnar et al., 2019; Rakic, 2009; Sun & Hevner, 2014). In this context, identifying the features that characterize the development specifically of the human neocortex is a fundamental challenge. Towards this goal, comparing the development of the human neocortex with that of the chimpanzee, our closest living relative, holds great promise. However, while tissue of developing human neocortex can, in principle, be obtained and subjected to experimental studies, this is not the case for tissue of developing chimpanzee neocortex.

Thanks to the seminal work of a few laboratories (Kadoshima et al., 2013; Karzbrun et al., 2018; Lancaster et al., 2013; Pasca et al., 2015; Qian et al., 2016; Quadrato et al., 2017), the brain organoid technology provides a way out of this dilemma. A specific subtype of brain organoids, the cerebral organoids, are relatively small (a few mm in diameter) three-dimensional (3D) structured cell assemblies that can be grown from embryonic stem cells (ESCs) (in the case of human) or induced pluripotent stem cells (iPSCs) (in the case of human and chimpanzee) and that emulate cerebral tissue (Arlotta, 2018; Di Lullo & Kriegstein, 2017; Fischer et al., 2019; Heide et al., 2018; Kelava & Lancaster, 2016; Lancaster et al., 2013). Cerebral organoids have been shown to exhibit several (albeit not all) of the hallmarks of developing neocortical tissue, including the two principal germinal zones, the ventricular zone (VZ) and the subventricular zone (SVZ), as well as the two major classes of progenitor cells therein, the apical progenitors (APs) and the basal progenitors (BPs) (Heide et al., 2018; Kadoshima et al., 2013; Lancaster et al., 2013; Qian et al., 2016; Quadrato et al., 2017). The generation of the various types of cortical neurons from these progenitor cells has also been established for brain and cerebral organoids (Heide et al., 2018; Kadoshima et al., 2013; Lancaster et al., 2017; Lancaster et al., 2013; Qian et al., 2016; Quadrato et al., 2017; Velasco et al., 2019). Moreover, in the case of human cerebral organoids, it has been shown that these recapitulate gene expression programs of fetal human neocortex development (Bhaduri et al., 2020; Camp et al., 2015; Velasco et al., 2019). In light of these findings, cerebral organoids have emerged as a promising primate model system to study cortical neurogenesis and, by comparison between human cerebral organoids and cerebral organoids from non-human primates including chimpanzee, to search for crucial differences in this complex process that underlie the evolution of human-specific features of neocortical development (Heide et al., 2018; Kanton et al., 2019; Mora-Bermudez et al., 2016; Otani et al., 2016; Pollen et al., 2019).

One intriguing feature of human neocortical development pertains to the size of, and the number of neurons in, the neocortex, both of which are greater than in any other primate. This increase is thought to reflect a greater proliferative capacity of the cortical stem and progenitor cells (collectively referred to as cortical neural progenitor cells (cNPCs)) in human as compared to non-human primates, which ultimately results in the greater neocortical size and neuron number (Dehay et al., 2015; Fish et al., 2008; Florio & Huttner, 2014; Lui et al., 2011; Sun & Hevner, 2014). Livesey and colleagues were the first to compare cortical neurogenesis in cerebral organoids generated from macaque, chimpanzee and human iPSCs and demonstrated that differences in cortical neurogenesis between human and non-human primates can indeed be revealed by this technology (Otani et al., 2016). In addition, an independent study comparing human and chimpanzee iPSC-derived cerebral organoids uncovered a novel difference in AP mitosis between human and this non-human primate (Mora-Bermudez et al., 2016).

These and other studies (Kanton et al., 2019; Pollen et al., 2019) have established that the intrinsic behaviour of cNPCs and the neuron generation therefrom as observed in human vs. chimpanzee cerebral organoids allows the identification of human-specific features of neocortical development. However, cerebral organoids also offer the opportunity of extrinsic genetic manipulation (Fischer et al., 2019). This is particularly relevant in the case of human-specific genes that in developing neocortex are preferentially expressed in cNPCs and hence have been implicated in human-specific features of neocortical development (Fiddes et al., 2018; Florio et al., 2015; Florio et al., 2018; Suzuki et al., 2018). Thus, to date, a human–chimpanzee cerebral organoid comparison to explore whether such human-specific genes are responsible for a human-type cNPC proliferative capacity has not yet been carried out. Examining such human-specific genes for their function in, and effects on, cNPC proliferation in cerebral organoids of human and chimpanzee, respectively, could not only provide corroborating evidence in support of their presumptive role in neocortical development during human evolution, but also open up new avenues to dissect their mechanism of action.

*ARHGAP11B* is a human-specific gene (Dennis et al., 2017; Sudmant et al., 2010) and the first such gene to have been implicated in human neocortical development and evolution (Florio et al., 2015; Florio et al., 2016; Heide et al., 2020; Kalebic et al., 2018). In fetal human neocortex, *ARHGAP11B* is preferentially expressed in cNPCs (Florio et al., 2015; Florio et al., 2018). When overexpressed in embryonic mouse and ferret neocortex, *ARHGAP11B* has been found to increase the proliferation and abundance of BPs (Florio et al., 2015; Kalebic et al., 2018), the cNPC class implicated in neocortical expansion during human development and evolution (Borrell & Götz, 2014; Dehay et al., 2015; Florio & Huttner, 2014; Lui et al., 2011). Moreover, a very recent study in which *ARHGAP11B* was expressed under the control of its own promoter to physiological levels in the fetal neocortex of the common marmoset has demonstrated that this human-specific gene can indeed induce the hallmarks of neocortical expansion in this non-human primate, increasing neocortex size, folding, BP levels and upper-layer neuron numbers (Heide et al., 2020). This study therefore established that *ARHGAP11B* is sufficient to expand primate BPs.

The ability of *ARHGAP11B* to increase the proliferation and abundance of BPs has been attributed not to the gene as it arose ≈5 mya by partial duplication of the widespread gene *ARHGAP11A* (Dennis et al., 2017; Sudmant et al., 2010), referred to as ancestral *ARHGAP11B*, but to an *ARHGAP11B* gene that subsequently underwent a point mutation, referred to as modern *ARHGAP11B* (Florio et al., 2016). Because of this point mutation, modern *ARHGAP11B* encodes a protein that contains a novel, human-specific C-terminal protein sequence (Florio et al., 2015; Florio et al., 2016) and that – in contrast to the nuclear ARHGAP11A protein – is imported into mitochondria where it expands BPs by promoting glutaminolysis, an effect involving its human-specific C-terminal protein sequence (Namba et al., 2020).

In addition to *ARHGAP11B*, at least 14 other human-specific genes with preferential expression in cNPCs have been identified (Florio et al., 2018). One of these, *NOTCH2NL*, has been shown, like *ARHGAP11B*, to increase the abundance of cycling BPs upon overexpression in embryonic mouse neocortex (Fiddes et al., 2018; Florio et al., 2018; Suzuki et al., 2018). However, in contrast to the ARHGAP11B protein, the NOTCH2NL protein is not localized in mitochondria and accordingly increases the abundance of cycling BPs via a distinct pathway (Fiddes et al., 2018; Suzuki et al., 2018). Considering these sets of findings together, the question arises to which extent *ARHGAP11B* contributes to the increase in cycling BPs in the context of the expansion of the neocortex in the course of human evolution. A first clue in this regard was obtained by the observation that a truncated form of the ARHGAP11A protein, ARHGAP11A220, which acts in a dominant-negative manner on ARHGAP11B’s ability to amplify BPs in embryonic mouse neocortex, reduces the abundance of cycling BPs in fetal human neocortical tissue ex vivo (Namba et al., 2020). Although this finding would be consistent with the notion that maintenance of the full level of cycling BPs in fetal human neocortex involves a contribution by ARHGAP11B, it remains to be established that ARHGAP11B is the only target of ARHGAP11A220, i.e. that the reduction in cycling BP abundance in fetal human neocortical tissue upon ARHGAP11A220 expression (Namba et al., 2020) was due to its interference with the action of ARHGAP11B rather than another target protein.

Hence, a key question regarding ARHGAP11B’s role in fetal human neocortical development is: Is ARHGAP11B required to maintain the full level of BP proliferation and abundance in human cerebral organoids? And conversely: Can the human-specific *ARHGAP11B* gene increase the proliferation and abundance of BPs when expressed in cerebral organoids of the chimpanzee, our closest living relative? In the present study, we have addressed these questions. In doing so, we provide support for the notion that the reduction in cycling BP abundance by ARHGAP11A220 indeed reflects its specific interference with the action of ARHGAP11B rather than another target protein. Importantly, we find that *ARHGAP11B* is necessary to maintain in human cerebral organoids, and sufficient to increase in chimpanzee cerebral organoids, BP proliferation and abundance, providing direct evidence in support of an indispensable role of ARHGAP11B in neocortical expansion during human development and evolution.

## Results

### Genetic manipulation of human and chimpanzee cerebral organoids by transfection of Aps

To obtain the data presented in this study, human and chimpanzee cerebral organoids were grown from human iPSCs of the line SC102A1 (Camp et al., 2015; Kanton et al., 2019; Mora-Bermudez et al., 2016) and chimpanzee iPSCs of the line Sandra A (Kanton et al., 2019; Mora-Bermudez et al., 2016), respectively. Cerebral organoid growth was carried out for 51-55 days according to an established protocol (Camp et al., 2015; Kanton et al., 2019; Lancaster & Knoblich, 2014; Lancaster et al., 2013; Mora-Bermudez et al., 2016), which involves the generation of embryoid bodies followed by their transformation into 3D cerebral tissue exhibiting numerous ventricular structures (Figure 1–figure supplement 1A). Various mixtures of DNA constructs, consisting of a cytosolic-GFP expression vector and either an expression vector with the cDNA of interest or the corresponding control vector, were then microinjected into the lumen of the larger ventricle-like structures within the cerebral organoids, followed by electroporation to transfect the cNPCs in the VZ (Figure 1–figure supplement 1A, (Fischer et al., 2019; Lancaster et al., 2013; Li et al., 2017)). Depending on the specific scientific question asked, cerebral organoids were fixed 2-10 days after electroporation, in the case of 2 days with addition of BrdU 1 h prior to fixation as indicated (Figure 1–figure supplement 1A). Fixed cerebral organoids were subjected to immunohistochemical analyses, using GFP immunofluorescence to identify the targeted cNPCs and their progeny (Figure 1–figure supplement 1A).

**Figure 1.**
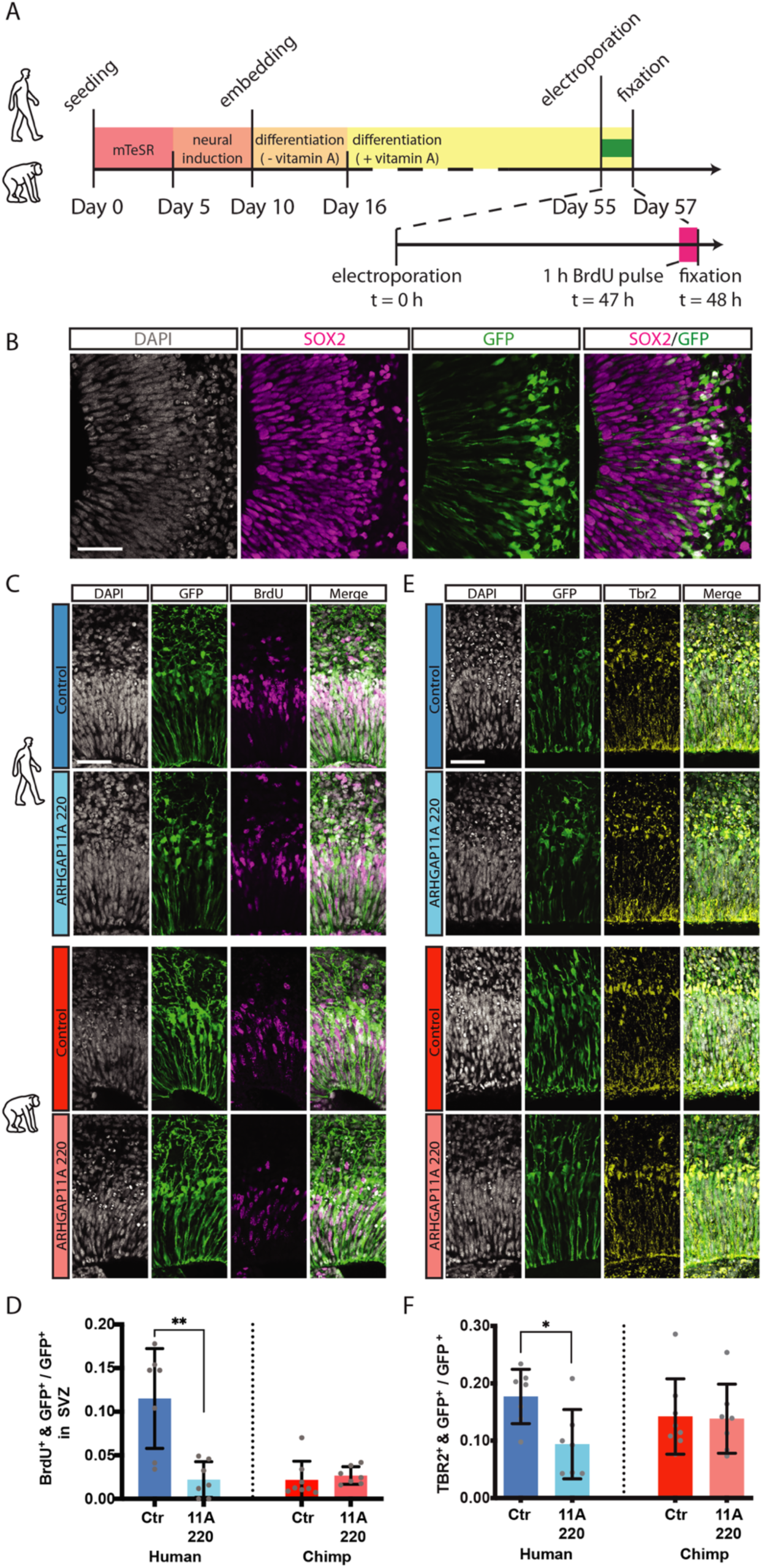
The dominant-negative effect of ARHGAP11A220 on ARHGAP11B’s ability to amplify BPs is specific and results in a marked reduction of BP proliferation and abundance in human cerebral organoids. (**A**) Timeline of human and chimpanzee cerebral organoid production with electroporation at day 55 and fixation 48 h later (day 57), with BrdU labeling 1 h prior to fixation. (**B**) Double immunofluorescence for SOX2 (magenta) and GFP (green), combined with DAPI staining (white), of a 57 days-old chimpanzee cerebral organoid 48 h after electroporation with GFP and control plasmids. Scale bar, 50 µm. (**C**) Double immunofluorescence for GFP (green) and BrdU (magenta), combined with DAPI staining (white), of 57 days-old human (upper panels) and chimpanzee (lower panels) cerebral organoids 48 h after electroporation with either control or ARHGAP11A220 plasmids as indicated. Scale bar, 50 µm. (**D**) Quantification of the proportion of GFP^+^ cells in the SVZ that are BrdU^+^ in 57 days-old human (blue) and chimpanzee (red) cerebral organoids 48 h after electroporation with either control (dark blue and dark red) or ARHGAP11A220 (light blue and light red) plasmids. Data are the mean of 7 human control, 7 human ARHGAP11A220, 8 chimpanzee control and 7 chimpanzee ARHGAP11A220 cerebral organoids; error bars indicate SD; **P<0.01. (**E**) Double immunofluorescence for GFP (green) and TBR2 (yellow), combined with DAPI staining (white), of 57 days-old human (upper panels) and chimpanzee (lower panels) cerebral organoids 48 h after electroporation with either control or ARHGAP11A220 plasmids as indicated. Scale bar, 50 µm. (**F**) Quantification of the proportion of GFP^+^ cells that are TBR2^+^ in 57 days-old human (blue) and chimpanzee (red) cerebral organoids 48 h after electroporation with either control (dark blue and dark red) or ARHGAP11A220 (light blue and light red) plasmids. Data are the mean of 7 human control, 7 human ARHGAP11A220, 8 chimpanzee control and 7 chimpanzee ARHGAP11A220 cerebral organoids; error bars indicate SD; *P<0.05.

A representative example of a control vector-transfected chimpanzee cerebral organoid 2 days after electroporation is presented in Figure 1–figure supplement 1B, showing that the majority of the GFP-positive cells were still observed in the VZ, co-localizing with the marker of proliferating cNPCs, SOX2. These data are consistent with the length of the total cell cycle of APs observed in chimpanzee cerebral organoids of ≈2 days (Mora-Bermudez et al., 2016) and suggest that the GFP-positive cells observed in the VZ 2 days after electroporation were either targeted APs, daughter APs of targeted APs, or newborn BPs derived from targeted APs.

### Requirement of ARHGAP11B for a human-type level of cycling BPs in human cerebral organoids

We first examined the role of ARHGAP11B on BP proliferation and abundance in human cerebral organoids. To this end, we made use of a truncated form of the ARHGAP11A protein (ARHGAP11A220) that has previously been shown to act in a dominant-negative manner on ARHGAP11B’s ability to amplify BPs (Namba et al., 2020). This dominant-negative action can be explained by the findings that ARHGAP11A220, via its truncated GAP domain, can interact with the same downstream effector system as ARHGAP11B, however without being able to change its activity, which requires the human-specific C-terminal domain of ARHGAP11B (Namba et al., 2020). To examine the effects of ARHGAP11A220 in human cerebral organoids, we used the same experimental protocol as described for Figure 1–figure supplement 1A, with a 2-day period between electroporation and analysis and a 1 h BrdU pulse prior to fixation (Figure 1A). SOX2 immunostaining was used to distinguish cNPCs in the VZ vs. SVZ and to identify the location of the GFP-positive progeny relative to these two germinal zones (Figure 1B, control electroporation). We found that compared to control, transfection of the cNPCs in the VZ of human cerebral organoids with the dominant-negative ARHGAP11A220 resulted in a marked reduction, down to the level observed in chimpanzee cerebral organoids, in the proportion of the GFP-positive progeny of the targeted APs found in the SVZ that had incorporated BrdU (Figure 1C, D). These data would be consistent with ARHGAP11B being required to maintain a human-type level of proliferating BPs in human cerebral organoids.

To corroborate this conclusion, we analyzed the transfected human cerebral organoids for the occurrence of TBR2, a marker of BPs (Englund et al., 2005; Sessa et al., 2008) that in fetal human neocortex is typically expressed in the basal intermediate progenitor (bIP) subpopulation of BPs (Hevner, 2019; Mihalas et al., 2016). Transfection of the human cerebral organoids with ARHGAP11A220 caused a reduction down to 50% of control in the proportion of the GFP-positive progeny of the targeted APs that were TBR2-positive (Figure 1E, F). Hence, taken together, these data indicate that indeed, ARHGAP11B is required to maintain a human-type level of proliferating BPs in human cerebral organoids.

### Specificity of ARHGAP11A220’s dominant-negative effect on ARHGAP11B’s ability to amplify BPs

We performed the same type of experiment and analyses with chimpanzee cerebral organoids, which lack ARHGAP11B, to determine whether the effects of ARHGAP11A220 were specific for ARHGAP11B. Indeed, upon transfection of the cNPCs in the VZ of chimpanzee cerebral organoids with ARHGAP11A220 vs. control, we observed no change in the proportion of the GFP-positive progeny of the targeted APs that had incorporated BrdU (progeny in SVZ; Figure 1C, D) and that were TBR2-positive (Figure 1E, F). Hence, the reduction in the level of proliferating BPs observed upon transfection of human cerebral organoids with the dominant-negative ARHGAP11A220 reflected a specific effect on ARHGAP11B’s ability to amplify BPs.

### Increased cycling BP abundance upon expression of human-specific *ARHGAP11B* in chimpanzee cerebral organoids

We next investigated whether *ARHGAP11B* would increase BP proliferation and abundance when expressed in chimpanzee cerebral organoids. Again, we used the same experimental protocol as described for Figure 1–figure supplement 1A, with a 2-day period between electroporation and analysis and a 1 h BrdU pulse prior to fixation (Figure 2A). Also, SOX2 immunostaining was used to distinguish cNPCs in the VZ vs. SVZ and to identify the location of the GFP-positive progeny relative to these two germinal zones (Figure 2B, control electroporation). We found that compared to control, transfection of the cNPCs in the VZ of chimpanzee cerebral organoids with an *ARHGAP11B*-expressing construct did not result in a statistically significant increase in the proportion of the GFP-positive progeny of the targeted APs found in the SVZ that had incorporated BrdU (Figure 2C, E). This indicated that this 2-day period was not sufficient for ARHGAP11B to increase the abundance of BPs that had progressed to S-phase. In contrast, analysis of the transfected chimpanzee cerebral organoids by TBR2 immunofluorescence revealed a marked, two-fold increase in the proportion of the GFP-positive progeny of the targeted APs that were TBR2-positive (Figure 2D, F). These data did point to an ability of ARHGAP11B to increase the generation of BPs in chimpanzee cerebral organoids, even if these cNPCs had not yet reached S-phase.

**Figure 2.**
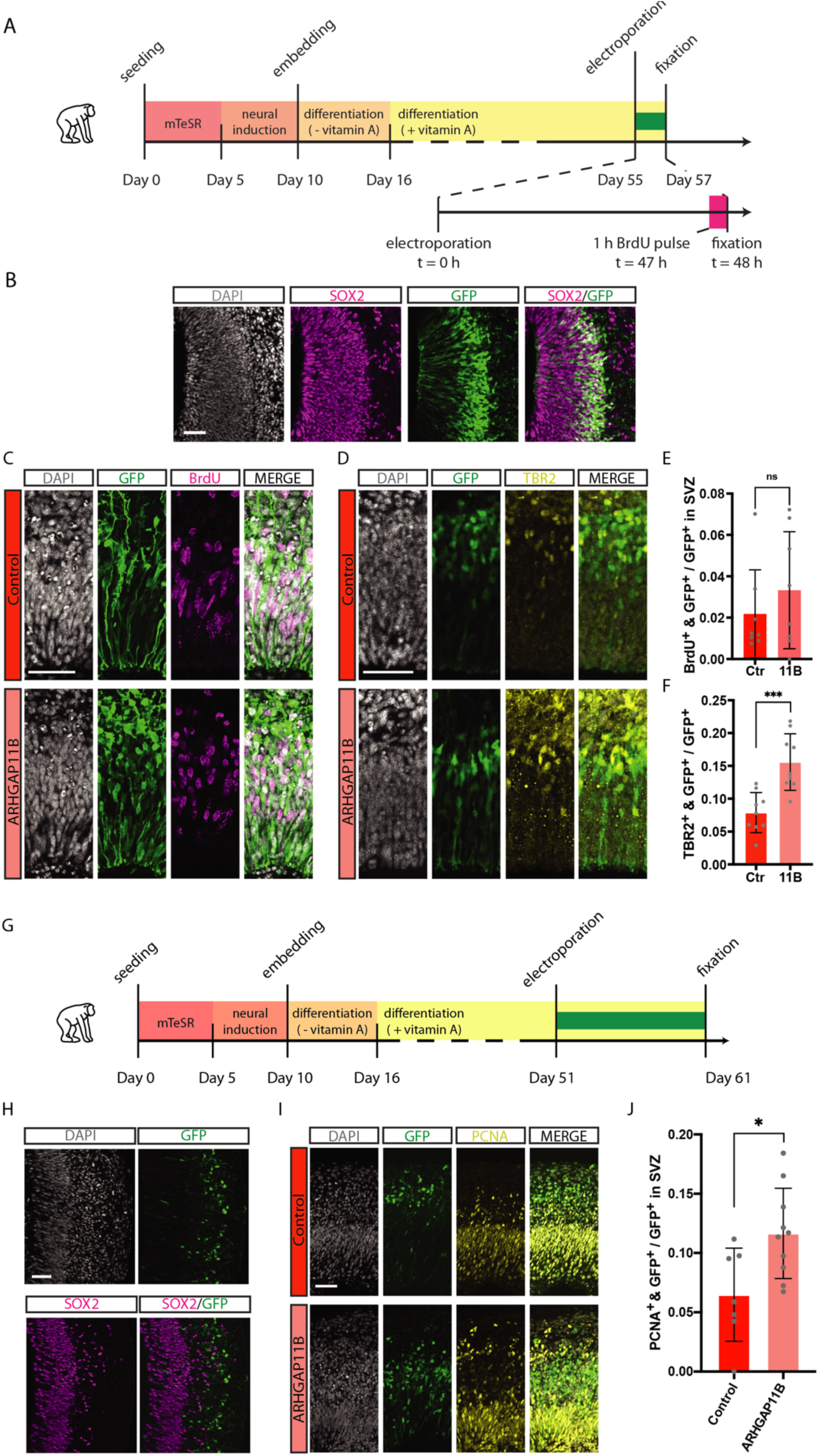
Expression of *ARHGAP11B* in chimpanzee cerebral organoids increases the abundance of cycling BPs. (**A**) Timeline of chimpanzee cerebral organoid production with electroporation at day 55 and fixation 48 h later (day 57), with BrdU labeling 1 h prior to fixation. (**B**) Double immunofluorescence for SOX2 (magenta) and GFP (green), combined with DAPI staining (white), of a 57 days-old chimpanzee cerebral organoid 48 h after electroporation with GFP and control plasmids. Scale bar, 50 µm. (**C**) Double immunofluorescence for GFP (green) and BrdU (magenta), combined with DAPI staining (white), of a 57 days-old chimpanzee cerebral organoid 48 h after electroporation with either control (top) or *ARHGAP11B* (bottom) plasmids. Scale bar, 50 µm. (**D**) Double immunofluorescence for GFP (green) and TBR2 (yellow), combined with DAPI staining (white), of a 57 days-old chimpanzee cerebral organoid 48 h after electroporation with either control (top) or *ARHGAP11B* (bottom) plasmids. Scale bar, 50 µm. (**E**) Quantification of the proportion of GFP^+^ cells in the SVZ that are BrdU^+^ in 57 days-old chimpanzee cerebral organoids 48 h after electroporation with either control (dark red) or *ARHGAP11B* (light red) plasmids. Data are the mean of 8 control and 8 ARHGAP11B cerebral organoids; error bars indicate SD; ns, not significant. (**F**) Quantification of the proportion of GFP^+^ cells that are TBR2^+^ in 57 days-old chimpanzee cerebral organoids 48 h after electroporation with either control (dark red) or *ARHGAP11B* (light red) plasmids. Data are the mean of 9 control and 9 ARHGAP11B cerebral organoids; error bars indicate SD; ***P<0.001. (**G**) Timeline of chimpanzee cerebral organoid production with electroporation at day 51 and fixation 10 days later (day 61). (**H**) Double immunofluorescence for SOX2 (magenta) and GFP (green), combined with DAPI staining (white), of a 61 days-old chimpanzee cerebral organoid 10 days after electroporation with GFP and control plasmids. Scale bar, 50 µm. (**I**) Double immunofluorescence for GFP (green) and PCNA (yellow), combined with DAPI staining (white), of a 61 days-old chimpanzee cerebral organoid 10 days after electroporation with either control (top) or *ARHGAP11B* (bottom) plasmids. Scale bar, 50 µm. (**J**) Quantification of the proportion of GFP^+^ cells in the SVZ that are PCNA^+^ in 61 days-old chimpanzee cerebral organoids 10 days after electroporation with either control (dark red) or *ARHGAP11B* (light red) plasmids. Data are the mean of 7 control and 10 ARHGAP11B cerebral organoids; error bars indicate SD; *P<0.05.

To further analyze the effect of *ARHGAP11B* on BP abundance in chimpanzee cerebral organoids, we used the same experimental protocol as described for Figure 1–figure supplement 1A, with a 10-day period between electroporation and analysis (Figure 2G). This longer period should allow the targeted APs to carry out multiple rounds of BP-generating cell divisions, thereby increasing the proportion of BPs among the GFP-positive progeny of the targeted APs. Accordingly, immunohistochemistry of chimpanzee cerebral organoids 10-days after electroporation for GFP and SOX2 revealed that the majority of the GFP-positive cells were observed in regions basal to the VZ, co-localizing less with the proliferating cNPC marker SOX2 in the VZ (Figure 2H, control electroporation). Compared to control, transfection of the chimpanzee cerebral organoids with *ARHGAP11B* caused a doubling in the proportion of the GFP-positive progeny of the targeted APs in the SVZ that were PCNA-positive, that is, cycling BPs (Figure 2I, J).

Taken together, these results indicate that the human-specific gene *ARHGAP11B*, similar to results previously obtained in embryonic mouse (Florio et al., 2015), embryonic ferret (Kalebic et al., 2018) and fetal marmoset (Heide et al., 2020) neocortex, can substantially increase the abundance of cycling BPs in developing cerebral cortex-like tissue of our closest living relative, the chimpanzee.

### Reduced cortical neuron generation concomitant with increased cycling BP abundance in chimpanzee cerebral organoids upon *ARHGAP11B* expression

If the increase in the abundance of cycling BPs in chimpanzee cerebral organoids upon *ARHGAP11B* expression reflected an increased proliferation of BPs, that is, BPs dividing to generate more BPs rather than neurons, one would expect a concomitant reduction in the generation of neurons from these cNPCs. To explore this possibility, we again used the same experimental protocol as described for Figure 1–figure supplement 1A, with a 10-day period between electroporation and analysis (Figure 3A) as this had revealed the increase in the abundance of cycling BPs. Compared to control, expression of *ARHGAP11B* in chimpanzee cerebral organoids resulted in a reduction in the proportion of the GFP-positive progeny of the targeted APs that were positive for the neuron markers Hu (Figure 3B, D) and NeuN (Figure 3C, E). In line with this GFP-positive progeny being neurons, the majority of the Hu-positive (Figure 3B) and NeuN-positive (Figure 3C) cells were located basally to the SVZ. We conclude that the increased abundance of cycling BPs observed after a 10-day period of expression of *ARHGAP11B* in chimpanzee cerebral organoids reflects an increased generation of BPs from BPs, resulting in a reduced generation of cortical neurons from BPs during this time period.

**Figure 3.**
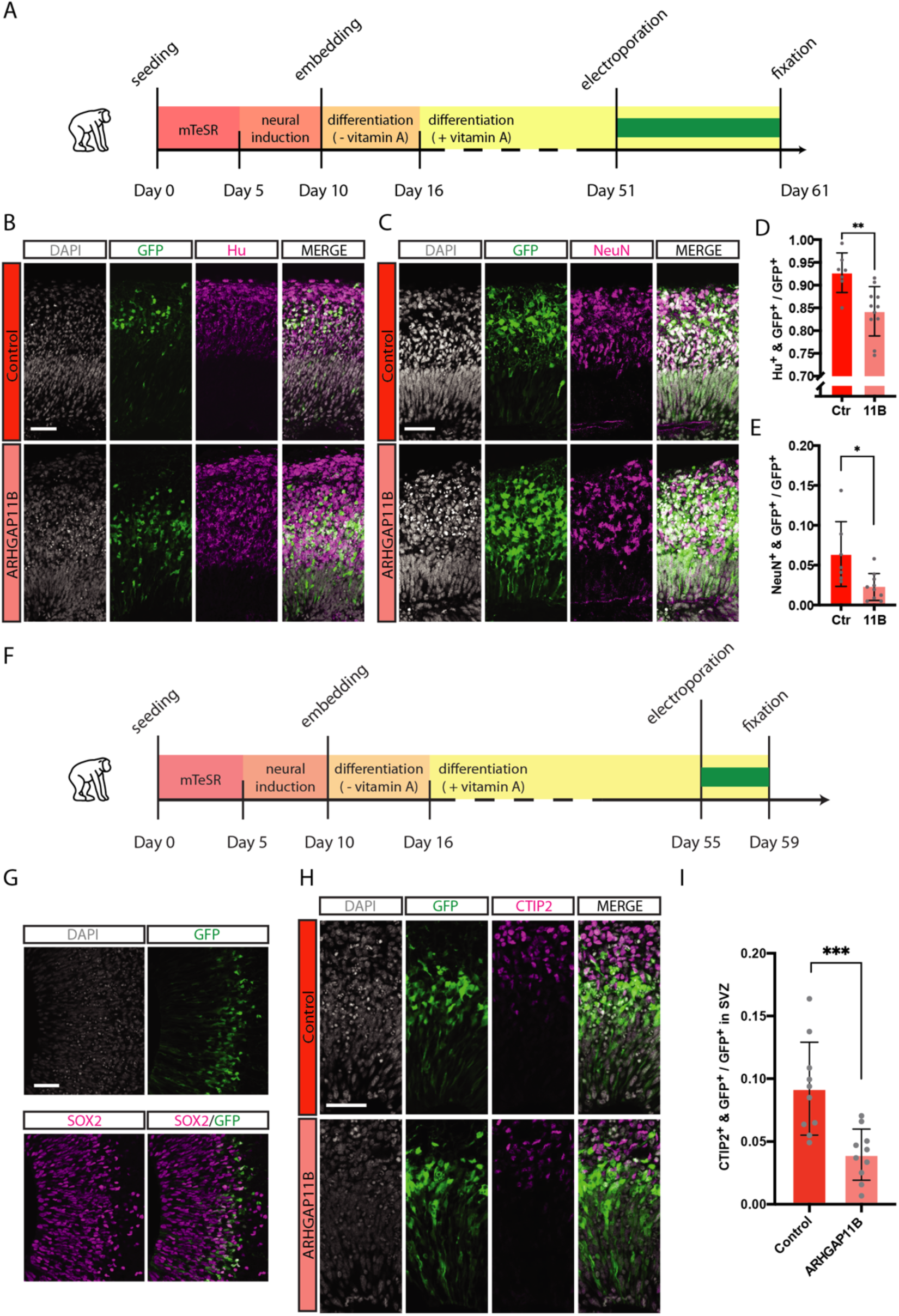
Expression of *ARHGAP11B* in chimpanzee cerebral organoids reduces the generation of cortical neurons. (**A**) Timeline of chimpanzee cerebral organoid production with electroporation at day 51 and fixation 10 days later (day 61). (**B**) Double immunofluorescence for GFP (green) and Hu (magenta), combined with DAPI staining (white), of a 61 days-old chimpanzee cerebral organoid 10 days after electroporation with either control (top) or *ARHGAP11B* (bottom) plasmids. Scale bar, 50 µm. (**C**) Double immunofluorescence for GFP (green) and NeuN (magenta), combined with DAPI staining (white), of a 61 days-old chimpanzee cerebral organoid 10 days after electroporation with either control (top) or *ARHGAP11B* (bottom) plasmids. Scale bar, 50 µm. (**D**) Quantification of the proportion of GFP^+^ cells that are Hu^+^ in 61 days-old chimpanzee cerebral organoids 10 days after electroporation with either control (dark red) or *ARHGAP11B* (light red) plasmids. Data are the mean of 7 control and 11 ARHGAP11B cerebral organoids; error bars indicate SD; **P<0.01. (**E**) Quantification of GFP^+^ cells that are NeuN^+^ in 61 days-old chimpanzee cerebral organoids 10 days after electroporation with either control (dark red) or *ARHGAP11B* (light red) plasmids. Data are the mean of 7 control and 10 ARHGAP11B cerebral organoids; error bars indicate SD; *P<0.05. (**F**) Timeline of chimpanzee cerebral organoid production with electroporation at day 55 and fixation four days later (day 59). (**G**) Double immunofluorescence for SOX2 (magenta) and GFP (green), combined with DAPI staining (white), of a 59 days-old chimpanzee cerebral organoid four days after electroporation with GFP and control plasmids. Scale bar, 50 µm. (**H**) Double immunofluorescence for GFP (green) and CTIP2 (magenta), combined with DAPI staining (white), of a 59 days-old chimpanzee cerebral organoid four days after electroporation with either control (top) or *ARHGAP11B* (bottom) plasmids. Scale bar, 50 µm. (**I**) Quantification of the proportion of GFP^+^ cells that are CTIP2^+^ in 59-days old chimpanzee cerebral organoids four days after electroporation with either control (dark red) or *ARHGAP11B* (light red) plasmids. Data are the mean of 10 control and 10 ARHGAP11B cerebral organoids; error bars indicate SD; ***P<0.001.

In the developing neocortex, the generation of cortical neurons begins with the production of deep-layer neurons, followed by the production of upper-layer neurons (Agirman et al., 2017; Cooper, 2008; Molyneaux et al., 2007). To investigate whether the first neurons generated in cerebral organoids would be of the deep-layer type, and whether this generation would be reduced upon *ARHGAP11B* expression, we used the same experimental protocol for chimpanzee cerebral organoids as described for Figure 1–figure supplement 1A, with a 4-day period between electroporation and analysis (Figure 3F). This time window should be just sufficient for two sequential rounds of cNPC division, (i) targeted APs generating GFP-positive BPs and (ii) GFP-positive BPs generating either GFP-positive neurons (control) or GFP-positive BPs (*ARHGAP11B*). Accordingly, SOX2 immunohistochemistry of chimpanzee cerebral organoids 4 days after electroporation revealed that the majority of the GFP-positive cells were observed in the basal VZ and in the SVZ (Figure 3G, control electroporation). Quantification of the deep-layer neuron marker CTIP2 (Arlotta et al., 2005; Molyneaux et al., 2007) indicated that compared to control, expression of *ARHGAP11B* in chimpanzee cerebral organoids resulted, after the 4-day period, in a reduction in the proportion of the GFP-positive progeny of the targeted APs that were CTIP2-positive (Figure 3H, I). This indicates that the first neurons generated by BPs in chimpanzee cerebral organoids, the generation of which is reduced upon *ARHGAP11B* expression due to the increased generation of BPs, are of the deep-layer type.

## Discussion

In conclusion, in the present study, the use of human and chimpanzee cerebral organoids has allowed us to investigate two key facets of the role of the human-specific gene *ARHGAP11B* in the evolutionary expansion of the human neocortex – whether it is necessary and whether it is sufficient for the increased abundance of cycling BPs that is thought to underlie this expansion. Each of the two cerebral organoid systems used here has unique advantages for addressing these questions. First, with regard to the human cerebral organoids, which have been shown to recapitulate many key features of fetal human neocortical tissue (Giandomenico et al., 2019; Heide et al., 2018; Kadoshima et al., 2013; Karzbrun et al., 2018; Lancaster et al., 2013; Qian et al., 2016; Quadrato et al., 2017), this system provides a readily available source of human neocortex-like tissue to investigate *ARHGAP11B*’s role during human neocortex development. Compared to fetal human neocortical tissue that can be obtained in principle, albeit only at an early stage of neocortex development, and studied ex vivo, human cerebral organoids offer a broader range of developmental stages. Moreover, because they originate from iPSCs, human cerebral organoids allow modes of genetic manipulation that are not possible with fetal human neocortical tissue ex vivo, such as the comprehensive ablation of a gene of interest. Also, studying the long-term effects of a manipulation, such as the effects of *ARHGAP11B* 10 days after electroporation into human cerebral organoids as done here, would be very difficult, if not impossible, with fetal human neocortical tissue ex vivo. In light of these advantages, we have used human cerebral organoids to investigate to which extent ARHGAP11B is necessary for the increased abundance of cycling BPs that is a characteristic of fetal human neocortex (Florio & Huttner, 2014; Lui et al., 2011). We find that interference with ARHGAP11B’s function results in a massive decrease in the level of cycling BPs, down to that observed in chimpanzee cerebral organoids. These data imply that ARHGAP11B is a major determinant of the increased abundance of cycling BPs in fetal human neocortex.

Second, the use of chimpanzee cerebral organoids has allowed us to determine *ARHGAP11B*’s role in neocortex expansion in the evolutionarily closest living species to human, and hence the contribution of this human-specific gene to neocortex expansion during primate evolution. Another human-specific gene, *NOTCH2NL*, has previously been studied in human and mouse brain organoids (Fiddes et al., 2018). This approach provided important insight into the function of NOTCH2NL and its potential role in neocortical expansion (Fiddes et al., 2018). However, to precisely determine the contribution of a human-specific gene to human neocortex expansion, it is necessary to study this gene in a model system that is evolutionarily as close as possible to humans. With regard to the human-specific gene *ARHGAP11B*, previous studies from our lab were performed in mouse (Florio et al., 2015; Florio et al., 2016), ferret (Kalebic et al., 2018) and the common marmoset (Heide et al., 2020), a non-human primate. However, considering the time point of origin of *ARHGAP11B* in the human lineage ≈5 mya, that is, shortly after the split from the lineage leading to chimpanzee and bonobo ≈7 mya, the marmoset is evolutionarily quite distant, as the split of the lineage leading to human from the lineage leading to marmoset happened ≈40 mya. While our previous expression of *ARHGAP11B* in fetal marmoset neocortex made the point that *ARHGAP11B* can expand the primate neocortex (Heide et al., 2020), the present expression of *ARHGAP11B* in chimpanzee cerebral organoids provides insight into the actual contribution of this human-specific gene to neocortex expansion during primate evolution. Thus, our finding that ARHGAP11B expression in chimpanzee cerebral organoids results in a doubling of cycling BP levels demonstrates that *ARHGAP11B* is a major contributor to the increased abundance of cycling BPs that is thought to underlie the evolutionary expansion of the human neocortex.

In summary, by using human and chimpanzee cerebral organoids, we have shown that *ARHGAP11B* is (i) necessary to maintain the human-type level of BP proliferation and abundance in human cerebral cortex tissue, and (ii) sufficient to increase the abundance of cycling BPs to a human-type level in chimpanzee cerebral cortex tissue. In line with an increase in BP proliferation, that is, in BP divisions that generate more BPs, upon *ARHGAP11B* expression, the generation of cortical neurons was found to be reduced, as indicated by quantification of the first class of cortical neurons produced, the deep-layer neurons. It is important to realize that this reflects an only transient reduction of cortical neuron generation, as *ARHGAP11B*’s function is to first increase the abundance of cycling BPs, which will eventually result in an increased generation of cortical neurons. Taken together, the effects of ARHGAP11B functional interference and *ARHGAP11B* ectopic expression in cerebral organoids of human and chimpanzee, respectively, provide evidence for *ARHGAP11B*’s essential role in increasing cycling BP abundance, and hence in human neocortex expansion, during primate evolution.

## Materials and Methods

### Cell culture and generation of cerebral organoids

Human SC102A-1 (System Bioscience) and chimpanzee Sandra A iPSC lines (Camp et al., 2015; Kanton et al., 2019; Mora-Bermudez et al., 2016) were cultivated using standard feeder-free conditions in mTeSR1 (StemCell Technologies) on Matrigel-(Corning) coated plates and differentiated into cerebral organoids using previously published protocols (Camp et al., 2015; Kanton et al., 2019; Lancaster & Knoblich, 2014; Lancaster et al., 2013; Mora-Bermudez et al., 2016) (Figure 1-3, Figure 1–figure supplement 1). Briefly, 10,000 cells per well were seeded into 96-well Ultra-low attachment plates (Corning) in mTeSR containing 50 µM Y27632 (StemCell Technologies). Medium was changed after 48 h to mTeSR without Y27632. On day 5 after seeding, medium was changed to neural induction medium (DMEM/F12 (Gibco) containing 1% N2 supplement (Gibco), 1% Glutamax supplement (Gibco), 1% MEM non-essential-amino-acids (Gibco) and 1 µg/ml heparin (Sigma-Aldrich)) and changed every other day. On day 10 after seeding, embryoid bodies were embedded in Matrigel and transferred to differentiation medium (1:1 DMEM/F12 (Gibco) / Neuralbasal (Gibco) containing 0.5% N2 supplement (Gibco), 0.025% Insulin solution (Sigma-Aldrich), 1% Glutamax supplement (Gibco), 0.5% MEM non-essential-amino-acids (Gibco), 1% B27 supplement (without vitamin A, Gibco), 1% penicillin-streptomycin and 0.00035% 2-mercaptoethanol (Merck)) to an orbital shaker. Medium was changed every other day and on day 16 after seeding switched to differentiation medium containing B27 supplement with vitamin A. Cerebral organoids were further cultured in this differentiation medium until fixation, with medium changes every three days and electroporation as indicated (see Figure 1-3, Figure 1–figure supplement 1).

### Electroporation of cerebral organoids

For cerebral organoid electroporation, organoids were placed in an electroporation chamber filled with pre-warmed mTeSR1 medium. Three to six ventricle-like structures per organoid were microinjected with a solution containing 0.1% Fast Green (Sigma) in sterile PBS, 500 ng/µl of either pCAGGS vector (control, (Florio et al., 2015)), pCAGGS-*ARHGAP11B* vector (Florio et al., 2015) or pCAGGS-*ARHGAP11A220* vector (Namba et al., 2020), in all cases together with 500 ng/µl pCAGGS-*EGFP* (Florio et al., 2015). The ventricle-like structures were microinjected with the Fast Green-containing solution until visibly filled. Electroporations were performed with five 50-msec pulses of 80 V at 1 sec intervals. Electroporated cerebral organoids were further cultured in differentiation medium containing vitamin A for the indicated time until fixation (see Figure 1-3, Figure 1–figure supplement 1), with medium changes every three days.

### BrdU labelling of cerebral organoids

To label cerebral organoid cells in S-Phase, a 1 h BrdU pulse was applied 47 h after electroporation by replacing the culture medium (differentiation medium containing vitamin A) with culture medium containing in addition 15 µM BrdU. Cerebral organoids were fixed 1 h later (see below).

### Fixation and cryosectioning of cerebral organoids

Cerebral organoids were fixed at the indicated time points (see Figure 1-3, Figure 1–figure supplement 1) in 4% paraformaldehyde in 120 mM phosphate buffer pH 7.4 for 2 h at 4°C. Fixed cerebral organoids were sequentially incubated in the phosphate buffer containing 15% sucrose and then 30% sucrose, each time overnight, at 4°C, embedded in Tissue-Tek OCT (Sakura), and frozen on dry ice. Cryosections of 20 µm thickness were cut and stored at –20°C until further use.

### Immunohistochemistry

Immunohistochemistry was performed as previously described (Mora-Bermudez et al., 2016). The following primary antibodies were used: BrdU (mouse monoclonal, EXBIO, 11-286-C100, RRID:AB_10732986, 1:300), Ctip2 (rat monoclonal, Abcam, ab18465, RRID:AB_2064130, 1:500), GFP (chicken polyclonal, Aves Labs, GFP-1020, RRID:AB_10000240, 1:500), Hu (mouse monoclonal, Thermo Fisher, A-21271, RRID:AB_221488, 1:200), NeuN (rabbit polyclonal, Abcam, ab104225, RRID:AB_10711153, 1:300), PCNA (rabbit polyclonal, Abcam, ab2426, RRID:AB_303062, 1:300; mouse monoclonal, Millipore, CBL407, RRID:AB_93501, 1:300), Sox2 (goat polyclonal, R+D Systems, AF2018, RRID:AB_355110, 1:150), Tbr2 (rabbit polyclonal, Abcam, ab23345, RRID:AB_778267, 1:500).

For all immunostainings, antigen retrieval was performed in 0.01 M sodium citrate buffer (pH 6.0) for 1 h at 70°C, prior to the overnight incubation with primary antibodies. The following secondary antibodies were used at a concentration of 1:500: Cy2: anti-chicken (donkey polyclonal, Dianova, 703-225-155, RRID:AB_2340370); Alexa Fluor 488: anti-chicken (goat polyclonal, Thermo Fisher, A-11039, RRID:AB_142924); Alexa Fluor 555: anti-goat (donkey polyclonal, Thermo Fisher, A-21432, RRID:AB_141788), anti-mouse (donkey polyclonal, Thermo Fisher, A-31570, RRID:AB_2536180), anti-rabbit (donkey polyclonal, Thermo Fisher, A-31572, RRID:AB_162543), anti-rat (goat polyclonal, Thermo Fisher, A-21434, RRID:AB_2535855); Alexa Fluor 594: anti-goat (donkey polyclonal, Thermo Fisher, A-11058, RRID:AB_2534105), anti-mouse (donkey polyclonal, Thermo Fisher, A-21203, RRID:AB_141633), anti-rabbit (donkey polyclonal, Thermo Fisher, A-21207, RRID:AB_141637); Alexa Fluor 647: anti-mouse (donkey polyclonal, Thermo Fisher, A-31571, RRID:AB_162542), anti-rabbit (donkey polyclonal, Thermo Fisher, A-31573, RRID:AB_2536183). All immunostained cryosections were counterstained with DAPI.

### Image Acquisition

Images were acquired using a Zeiss LSM 880 with 10x, 20x and 40x objectives. Images were taken as stacks of five 1-µm optical sections. When images were taken as tile scans, they were stitched together using the Zeiss ZEN software.

### Quantifications

All quantifications were performed blindly. Primary data were processed and results plotted using Prism (GraphPad Software). For all quantifications, cerebral organoids from two independent batches were used. Cell counts were performed in Fiji or Imaris. The mean of several electroporated ventricle-like structures was calculated. The data obtained from the quantifications are expressed as a proportion of the GFP-positive cell population.

### Statistical Analysis

All statistical analyses were conducted using Prism (GraphPad Software). Sample sizes (number of organoids per condition) are indicated in the figure legends. Student’s t-tests were used for statistical analyses. Statistical significances are indicated in the figure legends.

## Acknowledgments

We apologize to all researchers whose work could not be cited due to space limitations. We thank J. Peychl and his team of the Light Microscopy Facility at MPI-CBG for help with microscopy; C. Eugster and her team of the Organoid and Stem Cell Facility for organoid maintenance; and members of the Huttner laboratory for critical discussion. W.B.H. was supported by an ERA-NET NEURON (MicroKin) grant.

## Author contributions

Conceptualization: J.F., W.B.H and M.H.; Resources: T.N.; Investigation: J.F., J.P. and M.H.; Formal Analysis: J.F. and M.H.; Writing: J.F., W.B.H. and M.H.; Funding acquisition: W.B.H.; Supervision: W.B.H. and M.H.

## Competing interests

The authors declare no competing interests.

## Supplemental figures

**Figure 1–figure supplement 1.**
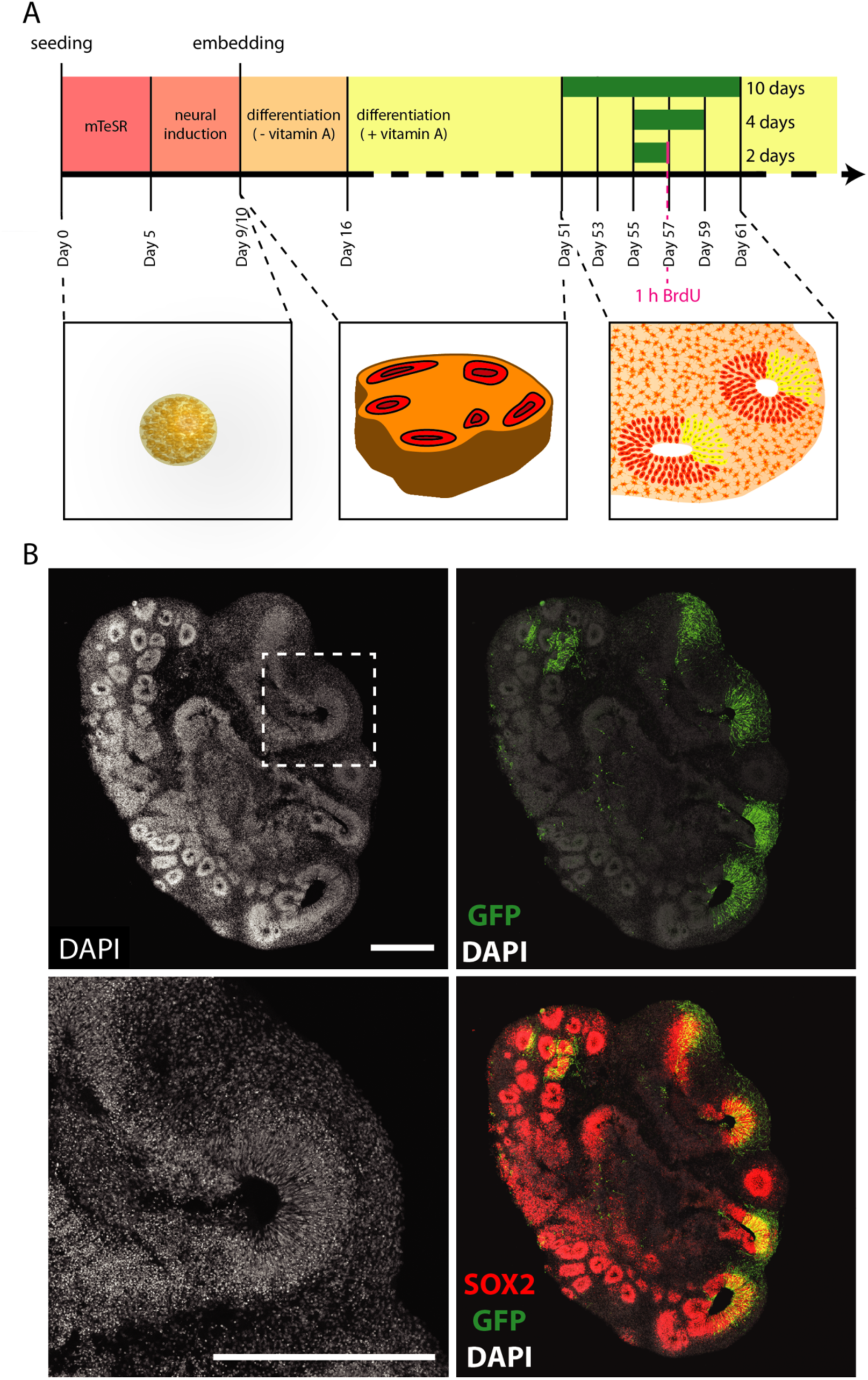
Experimental protocol of cerebral organoid production, electroporation and fixation. (**A**) Top: Timeline of cerebral organoid production detailing media as well as timepoints of electroporation, duration of vector expression (green bars; 2, 4 and 10 days) and timepoints of fixation for cerebral organoids. Bottom: Cartoons depicting embryoid body and cerebral organoids at various stages. Electroporated cells are indicated in yellow in the right image. (**B**) Double immunofluorescence for GFP (green) and SOX2 (red), combined with DAPI staining (white), of a 57 days-old chimpanzee cerebral organoid 48 h after electroporation with GFP and control plasmids. Scale bars, 500 µm.

## References

Agirman, G., Broix, L., & Nguyen, L. (2017). Cerebral cortex development: an outside-in perspective. FEBS Lett, 591(24), 3978–3992. https://doi.org/10.1002/1873-3468.12924

Arlotta, P. (2018). Organoids required! A new path to understanding human brain development and disease. Nat Methods, 15(1), 27–29. https://doi.org/10.1038/nmeth.4557

Arlotta, P., Molyneaux, B. J., Chen, J., Inoue, J., Kominami, R., & Macklis, J. D. (2005). Neuronal subtype-specific genes that control corticospinal motor neuron development in vivo. Neuron, 45(2), 207–221. https://doi.org/10.1016/j.neuron.2004.12.036

Bhaduri, A., Andrews, M. G., Mancia Leon, W., Jung, D., Shin, D., Allen, D., Jung, D., Schmunk, G., Haeussler, M., Salma, J., Pollen, A. A., Nowakowski, T. J., & Kriegstein, A. R. (2020). Cell stress in cortical organoids impairs molecular subtype specification. Nature, 578(7793), 142–148. https://doi.org/10.1038/s41586-020-1962-0

Borrell, V., & Götz, M. (2014). Role of radial glial cells in cerebral cortex folding. Curr. Opin. Neurobiol., 27, 39–46. https://doi.org/10.1016/j.conb.2014.02.007

Camp, J. G., Badsha, F., Florio, M., Kanton, S., Gerber, T., Wilsch-Bräuninger, M., Lewitus, E., Sykes, A., Hevers, W., Lancaster, M., Knoblich, J. A., Lachmann, R., Pääbo, S., Huttner, W. B., & Treutlein, B. (2015). Human cerebral organoids recapitulate gene expression programs of fetal neocortex development. Proc Natl Acad Sci U S A, 112(51), 15672–15677. https://doi.org/10.1073/pnas.1520760112

Cooper, J. A. (2008). A mechanism for inside-out lamination in the neocortex. Trends Neurosci, 31(3), 113–119. https://doi.org/10.1016/j.tins.2007.12.003

Silver, D. L., Grove, E. A., Haydar, T. F., Hensch, T. K., Huttner, W. B., Molnár, Z., Rubenstein, J. L., Sestan, N., Stryker, M. P., Sur, M., Tosches, M. A., & Walsh, C. A.. (2019). Evolution and ontogenetic development of cortical structures. In The Neocortex (Vol. 27, pp. 61–109). MIT Press.

Dehay, C., Kennedy, H., & Kosik, K. S. (2015). The outer subventricular zone and primate-specific cortical complexification. Neuron, 85(4), 683–694. https://doi.org/10.1016/j.neuron.2014.12.060

Dennis, M. Y., Harshman, L., Nelson, B. J., Penn, O., Cantsilieris, S., Huddleston, J., Antonacci, F., Penewit, K., Denman, L., Raja, A., Baker, C., Mark, K., Malig, M., Janke, N., Espinoza, C., Stessman, H. A. F., Nuttle, X., Hoekzema, K., Lindsay-Graves, T. A., Wilson, R. K., & Eichler, E. E. (2017). The evolution and population diversity of human-specific segmental duplications. Nat Ecol Evol, 1(3), 69. https://doi.org/10.1038/s41559-016-0069

Di Lullo, E., & Kriegstein, A. R. (2017). The use of brain organoids to investigate neural development and disease. Nat Rev Neurosci, 18(10), 573–584. https://doi.org/10.1038/nrn.2017.107

Englund, C., Fink, A., Lau, C., Pham, D., Daza, R. A., Bulfone, A., Kowalczyk, T., & Hevner, R. F. (2005). Pax6, Tbr2, and Tbr1 are expressed sequentially by radial glia, intermediate progenitor cells, and postmitotic neurons in developing neocortex. J. Neurosci., 25(1), 247–251. https://doi.org/10.1523/JNEUROSCI.2899-04.2005

Fiddes, I. T., Lodewijk, G. A., Mooring, M., Bosworth, C. M., Ewing, A. D., Mantalas, G. L., Novak, A. M., van den Bout, A., Bishara, A., Rosenkrantz, J. L., Lorig-Roach, R., Field, A. R., Haeussler, M., Russo, L., Bhaduri, A., Nowakowski, T. J., Pollen, A. A., Dougherty, M. L., Nuttle, X., Addor, M. C., Zwolinski, S., Katzman, S., Kriegstein, A., Eichler, E. E., Salama, S. R., Jacobs, F. M. J., & Haussler, D. (2018). Human-specific NOTCH2NL genes affect notch signaling and cortical neurogenesis. Cell, 173(6), 1356–1369. https://doi.org/10.1016/j.cell.2018.03.051

Fischer, J., Heide, M., & Huttner, W. B. (2019). Genetic modification of brain organoids. Front Cell Neurosci, 13, 558. https://doi.org/10.3389/fncel.2019.00558

Fish, J. L., Kennedy, H., Dehay, C., & Huttner, W. B. (2008). Making bigger brains -the evolution of neural-progenitor-cell division. J Cell Sci, 121, 2783–2793. https://doi.org/10.1242/jcs.023465

Florio, M., Albert, M., Taverna, E., Namba, T., Brandl, H., Lewitus, E., Haffner, C., Sykes, A., Wong, F. K., Peters, J., Guhr, E., Klemroth, S., Prüfer, K., Kelso, J., Naumann, R., Nüsslein, I., Dahl, A., Lachmann, R., Pääbo, S., & Huttner, W. B. (2015). Human-specific gene ARHGAP11B promotes basal progenitor amplification and neocortex expansion. Science, 347(6229), 1465–1470. https://doi.org/10.1126/science.aaa1975

Florio, M., Heide, M., Pinson, A., Brandl, H., Albert, M., Winkler, S., Wimberger, P., Huttner, W. B., & Hiller, M. (2018). Evolution and cell-type specificity of human-specific genes preferentially expressed in progenitors of fetal neocortex. eLife, 7, e32332. https://doi.org/10.7554/eLife.32332

Florio, M., & Huttner, W. B. (2014). Neural progenitors, neurogenesis and the evolution of the neocortex. Development, 141(11), 2182–2194. https://doi.org/10.1242/dev.090571

Florio, M., Namba, T., Pääbo, S., Hiller, M., & Huttner, W. B. (2016). A single splice site mutation in human-specific ARHGAP11B causes basal progenitor amplification. Sci Adv, 2(12), e1601941. https://doi.org/10.1126/sciadv.1601941

Giandomenico, S. L., Mierau, S. B., Gibbons, G. M., Wenger, L. M. D., Masullo, L., Sit, T., Sutcliffe, M., Boulanger, J., Tripodi, M., Derivery, E., Paulsen, O., Lakatos, A., & Lancaster, M. A. (2019). Cerebral organoids at the air-liquid interface generate diverse nerve tracts with functional output. Nat Neurosci, 22(4), 669–679. https://doi.org/10.1038/s41593-019-0350-2

Heide, M., Haffner, C., Murayama, A., Kurotaki, Y., Shinohara, H., Okano, H., Sasaki, E., & Huttner, W. B. (2020). Human-specific ARHGAP11B increases size and folding of primate neocortex in the fetal marmoset. Science, 369(6503), 546–550. https://doi.org/10.1126/science.abb2401

Heide, M., Huttner, W. B., & Mora-Bermudez, F. (2018). Brain organoids as models to study human neocortex development and evolution. Curr Opin Cell Biol, 55, 8–16. https://doi.org/10.1016/j.ceb.2018.06.006

Hevner, R. F. (2019). Intermediate progenitors and Tbr2 in cortical development. J Anat., 235, 616–625. https://doi.org/10.1111/joa.12939

Kadoshima, T., Sakaguchi, H., Nakano, T., Soen, M., Ando, S., Eiraku, M., & Sasai, Y. (2013). Self-organization of axial polarity, inside-out layer pattern, and species-specific progenitor dynamics in human ES cell-derived neocortex. Proc Natl Acad Sci U S A, 110(50), 20284–20289. https://doi.org/10.1073/pnas.1315710110

Kalebic, N., Gilardi, C., Albert, M., Namba, T., Long, K. R., Kostic, M., Langen, B., & Huttner, W. B. (2018). Human-specific ARHGAP11B induces hallmarks of neocortical expansion in developing ferret neocortex [Research Support, Non-U.S. Gov’t]. eLife, 7, e41241. https://doi.org/10.7554/eLife.41241

Kanton, S., Boyle, M. J., He, Z., Santel, M., Weigert, A., Sanchis-Calleja, F., Guijarro, P., Sidow, L., Fleck, J. S., Han, D., Qian, Z., Heide, M., Huttner, W. B., Khaitovich, P., Pääbo, S., Treutlein, B., & Camp, J. G. (2019). Organoid single-cell genomic atlas uncovers human-specific features of brain development. Nature, 574(7778), 418–422. https://doi.org/10.1038/s41586-019-1654-9

Karzbrun, E., Kshirsagar, A., Cohen, S. R., Hanna, J. H., & Reiner, O. (2018). Human brain organoids on a chip reveal the physics of folding. Nat Phys, 14(5), 515–522. https://doi.org/10.1038/s41567-018-0046-7

Kelava, I., & Lancaster, M. A. (2016). Stem cell models of human brain development. Cell Stem Cell, 18(6), 736–748. https://doi.org/10.1016/j.stem.2016.05.022

Lancaster, M. A., Corsini, N. S., Wolfinger, S., Gustafson, E. H., Phillips, A. W., Burkard, T. R., Otani, T., Livesey, F. J., & Knoblich, J. A. (2017). Guided self-organization and cortical plate formation in human brain organoids. Nat Biotechnol, 35(7), 659–666. https://doi.org/10.1038/nbt.3906

Lancaster, M. A., & Knoblich, J. A. (2014). Generation of cerebral organoids from human pluripotent stem cells. Nat. Protoc., 9(10), 2329–2340. https://doi.org/10.1038/nprot.2014.158

Lancaster, M. A., Renner, M., Martin, C. A., Wenzel, D., Bicknell, L. S., Hurles, M. E., Homfray, T., Penninger, J. M., Jackson, A. P., & Knoblich, J. A. (2013). Cerebral organoids model human brain development and microcephaly. Nature, 501(7467), 373–379. https://doi.org/ https://doi.org/10.1038/nature12517

Li, R., Sun, L., Fang, A., Li, P., Wu, Q., & Wang, X. (2017). Recapitulating cortical development with organoid culture in vitro and modeling abnormal spindle-like (ASPM related primary) microcephaly disease. Protein Cell, 8(11), 823–833. https://doi.org/10.1007/s13238-017-0479-2

Lui, J. H., Hansen, D. V., & Kriegstein, A. R. (2011). Development and evolution of the human neocortex. Cell, 146(1), 18–36. https://doi.org/10.1016/j.cell.2011.06.030

Mihalas, A. B., Elsen, G. E., Bedogni, F., Daza, R. A., Ramos-Laguna, K. A., Arnold, S. J., & Hevner, R. F. (2016). Intermediate progenitor cohorts differentially generate cortical layers and require Tbr2 for timely acquisition of neuronal subtype identity. Cell Rep, 16(1), 92–105. https://doi.org/10.1016/j.celrep.2016.05.072

Molnar, Z., Clowry, G. J., Sestan, N., Alzu’bi, A., Bakken, T., Hevner, R. F., Huppi, P. S., Kostovic, I., Rakic, P., Anton, E. S., Edwards, D., Garcez, P., Hoerder-Suabedissen, A., & Kriegstein, A. (2019). New insights into the development of the human cerebral cortex. J Anat, 235(3), 432–451. https://doi.org/10.1111/joa.13055

Molyneaux, B. J., Arlotta, P., Menezes, J. R., & Macklis, J. D. (2007). Neuronal subtype specification in the cerebral cortex. Nat. Rev. Neurosci., 8(6), 427–437. https://doi.org/nrn2151

Mora-Bermudez, F., Badsha, F., Kanton, S., Camp, J. G., Vernot, B., Kohler, K., Voigt, B., Okita, K., Maricic, T., He, Z., Lachmann, R., Pääbo, S., Treutlein, B., & Huttner, W. B. (2016). Differences and similarities between human and chimpanzee neural progenitors during cerebral cortex development. eLife, 5, e18683. https://doi.org/10.7554/eLife.18683

Namba, T., Doczi, J., Pinson, A., Xing, L., Kalebic, N., Wilsch-Bräuninger, M., Long, K. R., Vaid, S., Lauer, J., Bogdanova, A., Borgonovo, B., Shevchenko, A., Keller, P., Drechsel, D., Kurzchalia, T., Wimberger, P., Chinopoulos, C., & Huttner, W. B. (2020). Human-specific ARHGAP11B acts in mitochondria to expand neocortical progenitors by glutaminolysis. Neuron, 105(5), 867–881. https://doi.org/10.1016/j.neuron.2019.11.027

Otani, T., Marchetto, M. C., Gage, F. H., Simons, B. D., & Livesey, F. J. (2016). 2D and 3D stem cell models of primate cortical development identify species-specific differences in progenitor behavior contributing to brain size. Cell Stem Cell, 18(4), 467–480. https://doi.org/10.1016/j.stem.2016.03.003

Pasca, A. M., Sloan, S. A., Clarke, L. E., Tian, Y., Makinson, C. D., Huber, N., Kim, C. H., Park, J. Y., O’Rourke, N. A., Nguyen, K. D., Smith, S. J., Huguenard, J. R., Geschwind, D. H., Barres, B. A., & Pasca, S. P. (2015). Functional cortical neurons and astrocytes from human pluripotent stem cells in 3D culture. Nat Methods, 12(7), 671–678. https://doi.org/10.1038/nmeth.3415

Pollen, A. A., Bhaduri, A., Andrews, M. G., Nowakowski, T. J., Meyerson, O. S., Mostajo-Radji, M. A., Di Lullo, E., Alvarado, B., Bedolli, M., Dougherty, M. L., Fiddes, I. T., Kronenberg, Z. N., Shuga, J., Leyrat, A. A., West, J. A., Bershteyn, M., Lowe, C. B., Pavlovic, B. J., Salama, S. R., Haussler, D., Eichler, E. E., & Kriegstein, A. R. (2019). Establishing cerebral organoids as models of human-specific brain evolution. Cell, 176(4), 743–756. https://doi.org/10.1016/j.cell.2019.01.017

Qian, X., Nguyen, H. N., Song, M. M., Hadiono, C., Ogden, S. C., Hammack, C., Yao, B., Hamersky, G. R., Jacob, F., Zhong, C., Yoon, K. J., Jeang, W., Lin, L., Li, Y., Thakor, J., Berg, D. A., Zhang, C., Kang, E., Chickering, M., Nauen, D., Ho, C. Y., Wen, Z., Christian, K. M., Shi, P. Y., Maher, B. J., Wu, H., Jin, P., Tang, H., Song, H., & Ming, G. L. (2016). Brain-region-specific organoids using mini-bioreactors for modeling ZIKV exposure. Cell, 165(5), 1238–1254. https://doi.org/10.1016/j.cell.2016.04.032

Quadrato, G., Nguyen, T., Macosko, E. Z., Sherwood, J. L., Min Yang, S., Berger, D. R., Maria, N., Scholvin, J., Goldman, M., Kinney, J. P., Boyden, E. S., Lichtman, J. W., Williams, Z. M., McCarroll, S. A., & Arlotta, P. (2017). Cell diversity and network dynamics in photosensitive human brain organoids. Nature, 545(7652), 48–53. https://doi.org/10.1038/nature22047

Rakic, P. (2009). Evolution of the neocortex: a perspective from developmental biology. Nat. Rev. Neurosci., 10(10), 724–735. https://doi.org/nrn2719

Sessa, A., Mao, C. A., Hadjantonakis, A. K., Klein, W. H., & Broccoli, V. (2008). Tbr2 directs conversion of radial glia into basal precursors and guides neuronal amplification by indirect neurogenesis in the developing neocortex. Neuron, 60(1), 56–69. https://doi.org/S0896-6273(08)00807-6

Sudmant, P. H., Kitzman, J. O., Antonacci, F., Alkan, C., Malig, M., Tsalenko, A., Sampas, N., Bruhn, L., Shendure, J., & Eichler, E. E. (2010). Diversity of human copy number variation and multicopy genes. Science, 330(6004), 641–646. https://doi.org/10.1126/science.1197005

Sun, T., & Hevner, R. F. (2014). Growth and folding of the mammalian cerebral cortex: from molecules to malformations. Nat. Rev. Neurosci., 15(4), 217–232. https://doi.org/10.1038/nrn3707

Suzuki, I. K., Gacquer, D., Van Heurck, R., Kumar, D., Wojno, M., Bilheu, A., Herpoel, A., Lambert, N., Cheron, J., Polleux, F., Detours, V., & Vanderhaeghen, P. (2018). Human-specific NOTCH2NL genes expand cortical neurogenesis through Delta/Notch regulation. Cell, 173(6), 1370–1384. https://doi.org/10.1016/j.cell.2018.03.067

Velasco, S., Kedaigle, A. J., Simmons, S. K., Nash, A., Rocha, M., Quadrato, G., Paulsen, B., Nguyen, L., Adiconis, X., Regev, A., Levin, J. Z., & Arlotta, P. (2019). Individual brain organoids reproducibly form cell diversity of the human cerebral cortex. Nature, 570, 523–527. https://doi.org/10.1038/s41586-019-1289-x

